# Causality Mapping Using Resting-State fMRI reveals Suppressed Functional Connectivity in Schizophrenia Patients

**DOI:** 10.1101/2020.09.12.295048

**Authors:** Fayyaz Ahmed, Zunira Saghir, Namra Aamir, Turki Abualait, Safee Ullah Chaudhary, Shahid Bashir

## Abstract

Schizophrenia is a psychotic brain disorder in which patients exhibit aberrant connectivity between different regions of the brain. Neuroimaging is a state-of-the-art technique that is now increasingly been employed in clinical investigation of Schizophrenia. In the present study, we have used resting-state functional magnetic resonance neuroimaging (rsfMRI) to elucidate the cause-and-effect relationships among four regions of the brain including occipital, temporal, and frontal lobes and hippocampus in Schizophrenia. For that, we have employed independent component analysis, a seed-based temporal correlation analysis, and Granger causality analysis for measuring causal relationships amongst four regions of the brain in schizophrenia patients. Eighteen subjects with nine patients and nine controls were evaluated in the study. Our results show that Schizophrenia patients exhibit significantly different activation patterns across the selected regions of the brain in comparison with the control. In addition to that, we also observed an aberrant causal relationship between these four regions of the brain. In particular, the temporal and frontal lobes of patients with schizophrenia had a significantly lowered causal relationship with the other areas of the brain. Taken together, the study elucidates the dysregulated brain activity in Schizophrenia patients, decodes its causal mapping and provides novel insights towards employment in clinical evaluation of Schizophrenia.

## INTRODUCTION

Schizophrenia (SZ) is a chronic mental disorder affecting approximately 1% of the global population (1). The condition is characterized by cognitive and behavioral alterations, accompanied by hallucinations, delusions, and disorganized thinking (2,3). SZ typically develops in young adults between the age of 18 and 35 (4) and has a lifetime prevalence of 1% (4). Recent research on SZ has uncovered several genetic and environmental factors that correlates with the development of the disease (4). An increasing amount of experimental evidence is now suggesting that SZ induces extensive alterations in the connectivity between various regions of the brain that results in its functional dysregulation (5,6). Moreover, research studies have also reported significant reductions in functional connectivity amongst various regions of the brain in SZ patients (7–9). However, the type and scale of these disconnections, suppressed functional connectivity and the underlying pathophysiological mechanisms in SZ remain unclear (11).

Advancements in brain imaging techniques and protocols have now enabled scientists to carry out in-depth investigations of the neuropathological mechanisms of SZ (10–14). Several groundbreaking studies leveraging these technologies have helped reveal a wide variety of abnormal structural and functional connections among anatomically distant regions of the brain in SZ patients (15–18). Investigations using functional brain imaging have helped elucidate reduced activation in the frontal, striatal, and parietal regions during the performance of cognitive tasks in patients with SZ (19,20). Furthermore, altered patterns of neural activation have been reported in the prefrontal cortices, including the anterior cingulate cortex (ACC), the supplementary motor area (SMA), the pre-SMA, the parietal cortex, and the subcortical basal ganglia nuclei (20–23). Data from diffusion tensor imaging have highlighted reduced fractional anisotropy (FA) in the internal capsule, thalamus, corpus callosum (CC), white matter microstructures (24), and throughout the entire brain in patients with SZ (25). Reduced FA in the CC has been associated with higher lateral ventricle (LV) volume, while higher radial diffusivity (RD) values are associated with larger LV volume in patients with SZ (26). This suggests that SZ may involve abnormalities in structural white matter, which connects and activates different regions of the brain.

These findings have been supported and expanded upon by several neuroimaging studies on functional connectivity (FC), which have elucidated temporal associations between various regions of the brain (27). Specifically, this approach has been utilized to investigate disruptions in functional connectivity in brain of SZ patients (28). Previous studies have examined specific regions of the brain at resting state (29–31), and during sensory (32), and cognitive tasks (33) in patients with SZ. Reduced FC has been extensively reported in the neural pathways of patients with SZ, including the amygdala subregional-sensorimotor pathway (34), frontotemporal (35), thalamocortical (36), and cortico-cerebellar pathways (37). Differential FC in brain of patients with SZ and healthy controls, therefore, could have significant clinical implications, as psychopharmacological treatments could target aberrant FC in patients with SZ (39, 40). In addition, these differences could also be used as a neuroimaging biomarker to guide diagnoses of SZ (41, 42).

Towards this goal, in this work, we investigate the resting-state functional connectivity in brain of patients with SZ and examine the cause-and-effect relationships among these regions of the brain (38). For that, we have combined three different methods for analyzing FC, including (i) spatial independent component analysis (ICA) to identify the characteristics or features of fMRI data that are maximally independent, (ii) a seed-based temporal correlation analysis (SCA) to assess time-series associations and connections among regions, and (iii) Granger causality analysis (GCA) to analyze effective connectivity in the brain. In specific, spatial ICA is employed to extract the overall connectivity patterns at the whole-brain level and is used to elucidate the spatial structure of the blood-oxygen-level-dependent (BOLD) signal (47, 50, 51). The seed-based SCA was used to identify regions, or “seeds,” that correlate with FC and activity in other seeds or regions using time series of BOLD signals from seed-voxels and other regions (41–44). Lastly, the GCA was used to study the bidirectional effects between two variables in a time series (45) to examine time-lagged causal effects on specific regions of the brain by using time predictions between an fMRI time series (46,47). The approach helped identify the dynamic causal interactions between the four selected regions of the brain. Our results show that the brains of patients with SZ exhibit aberrant activation patterns in the HC, OL, TL, and FL. Moreover, the functional and structural connectivity among these four regions differs significantly in patients with SZ, which has been validated by the VAR models using Granger causality. Last, our causal analysis through mediation revealed significant differences in decision making in patients with SZ.

Taken together, this study expands our understanding of the resting-state FC (rsFC) in patients with SZ at the whole-brain level and reports the aberrations in FC among multiple regions of the brain using a combination of statistical approaches. The study also helps characterize the direction of FC among different regions of the brain and how it differs in patients with SZ from healthy controls (39) thereby providing valuable insights into the neural basis of SZ (39).

## METHODOLOGY

### 2.1 Granger causality

We used a Granger causality (G-causal) test and VAR modeling to analyze the fMRI data. This method determines the causal relationships between two variables. If a variable x has a G-causal relationship with another variable y, then the lag values of x can be used to forecast the future values of y and vice versa. This study examined the causal relationships among four regions of the brain: the frontal, occipital, and temporal lobes and the hippocampus.

#### Mathematical Formulation of Granger Causality

Consider two random variables Xt, Y_t_. Assume a lag length of p. For example, y is occipital lobe and X is temporal lobe. For occipital and temporal lobes, the model could be written as:

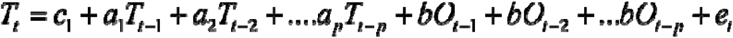

Estimate by OLS and test for the following hypothesis:

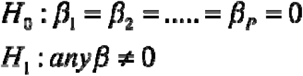

To check if occipital lobe does granger-cause to temporal lobe or not, here we will take y as a occipital lobe ‘o’ and x as a temporal lobe ‘t’.

Unrestricted sum of squared residual:

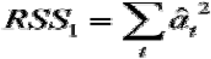

Restricted sum of square residual:

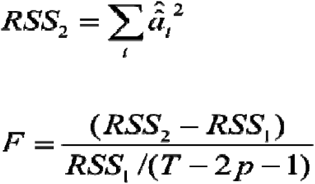

Under general condition, the OLS estimate is given by,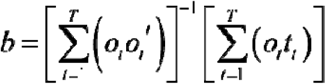 assuming that the (kxk) matrix 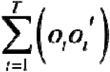 is nonsingular the OLS sample residual for observation t is 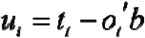.Often the model 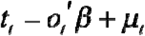 is written in matrix notation as

*T* − *Oβ* + *µ* Where

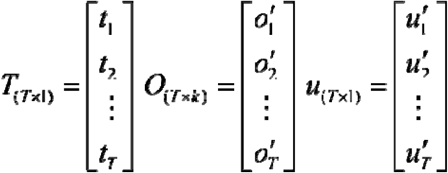

Then the OLS estimate in 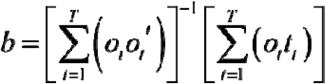 can be written as

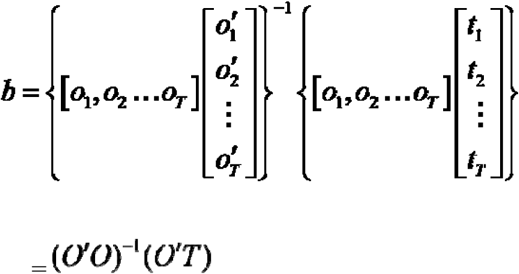

Similarly, residual can be written as:

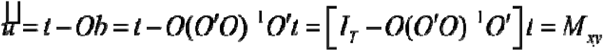

Where ***M***_***x***_ is defined as the following (T × T) matrix ***M***_***x***_=*I*_*T*_ – *O*(*O*’*O*)^−1^*O*’.One can readily verify that 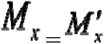 where idempotent ***M***_***x***_ ***M***_***x***_ ***= M***_***x***_ and the orthogonal to the columns of O. ***M***_***x***_X=0. OLS sample residuals are orthogonal to the explanatory variables in O and population residual can be found by substituting 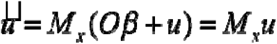.The difference between the OLS estimate b and the true population parameter β is found by

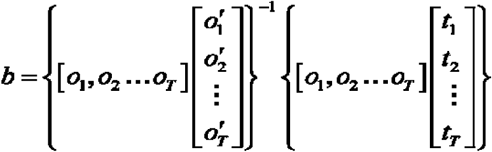

substituting *T*=*O β + μ* into

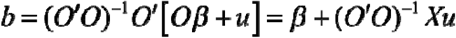

Now estimate the F test (Wald test) about beta under assumption. A Wald test of Ho is based on the following results. Consider an (nx1) vector z ∼ ***N*(0, Ω)** with non-singular then Z’**Ω** ^1^**Z** ∼ χ^2^(*n*)

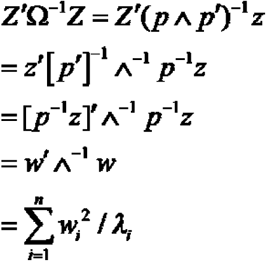

Where *w* = *p*^*–*1^*z* and w is Gaussian with mean zero and variance *E*(*ww*’) = *E*(*p*^−1^ *zz*’[*p*’]^−1^) = *p*^−1^Ω[*p*^−1^]^−1^ = *p*^−1^ *p* ^ *p*’ [*p’*]^−1^ = ^.Thus *w*’ ^^−1^ *w* is the sum of squares of n independent normal variables each divided by its variance *λ*.It accordingly has a χ^2^_*n*_ distribution as claimed. Applying proposition directly to *Rb* ∼ *N*(*r, δ*^2^*R*) (O’O)^−1^ *R*’) under Ho (*Rb* – *r*)’[*δ*^2^*R*(*O’O*)^−1^ *R’*]^−1^ (*Rb* – *r*) ∼ *χ*^2^_*m*_ Replacing σ with the estimate s and dividing by the number of restriction gives the Wald form of the OLS F test of a linear hypothesis. *F* = (*Rb* – *r*)’[*s*^2^ *R*(*O*’*O*) ^1^*R*’]^−1^ (*Rb* – *r*)/ *m*

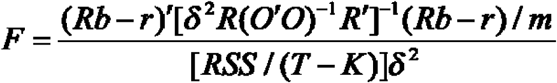

The numerator is a χ^2^_*m*_ variable divided by its degree of freedom while the denominator is χ^2^ (T-k) variable divided by its degree of freedom again since b and *μ* are independent, the numerator and denominator are independent of each other hence *F* – (*Rb* – *r*)’[*s*^2^*R*(*O*’*O*)^−1^ *R*^1^]^−1^ (*Rb – r*)/*m* has an exact F(m, T-K) distribution under Ho. Let b denote the unconstrained OLS estimate and let RSS_1_ be the residual sum of square resulting from using this estimate.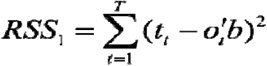 Let b* denote the constrained OLS estimate and RSS_o_ the residual sum squares from the constrained OLS estimation. 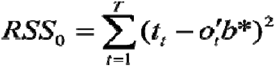. Then the Wald form of the F test of a linear hypothesis *F* = (*Rb* – *r*)’[*s*^2^*R*(*O*’*O*)^−1^ *R*^1^]^−1^ (*Rb – r*)/*m* can equivalently be calculated as 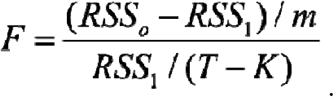.

### Vector auto regression (VAR) model

VAR models are used for estimating and forecasting in time series data. In multivariate time series analysis, it is one of most easy to use technique. It is known as extension of AR model. One of crucial step in VAR is selection of lags for lag selection we have three criteria’s AIC BIC HQIC.

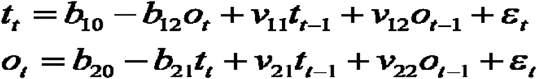

In above model ‘o’ represent the occipital lobe and ‘t’ represent the temporal lobe. VAR models are used for forecasting in time series but we can employ them to check the Granger causality of the variables. One of the important steps in VAR model is the selection of lag length which is based on specific criteria.

### Data

The study was approved by the Institutional Review Board of King Khalid University Hospital. The participants included a selection of 15 healthy controls and 15 patients with SZ. From these, we selected 18 subjects (9 controls and 9 patients with SZ). All participants underwent the same number of scans according to our study requirements. The participants were aged 33.14 ± 9.96 (mean ± SD) years. The SZ participants were recruited through local psychiatric clinics, and the controls were recruited from hospital volunteers. All participants provided informed written consent before participating. All SZ and control subjects were outpatients and had been clinically stable for at least two weeks prior to the study. The SZ participants were diagnosed by a research psychiatrist based on the DSM.IV criteria; the diagnoses were confirmed by a trained research assistant.

Written informed consent was obtained from subjects which was approved by the Institutional Review Board (IRB) at King Khalid University Hospital (KKUH). All procedures were conducted according to the Declaration of Helsinki.

## 3. RESULTS

### 3.1. The brains of patients with SZ exhibit aberrant activation patterns in the hippocampus (HC), occipital lobe (OL), temporal lobe (TL), and frontal lobe (FL)

To evaluate differences in brain activation patterns in patients with SZ, we obtained fMRI scans of nine subjects with SZ (**Supplementary Data 1**). Nine clinically healthy individuals were also scanned and used as controls (**Supplementary Data 2**). The resulting fMRI scan data were pre-processed for realignment, spatial normalization, smoothing, and co-registration using statistical parametric mapping (SPM12) (48) (**Figures 1A, B**). We then compared activation levels in four regions of interest (ROIs): the hippocampus (HC), the occipital lobe (OL), the temporal lobe (TL) and the frontal lobe (FL). Brain activation in patients with SZ was compared to that of the controls using two independent sample *t*-tests (49). We found that average brain activity in the two groups differed significantly in the HC (*t*(798) = 125.254, p < 0.05), the OL (*t* (798) = 43.573, p < 0.05), the TL (*t*(798) = 130.784, p < 0.05), and the FL (*t*(798) = −9.774, p < 0.05) (**Figure 1C**). We conclude that patients with SZ have significantly less activation in the HC, the OL, and the TL than healthy controls (59). However, we also observed significantly higher activation in the FL of patients with SZ.

**Figure 1.**
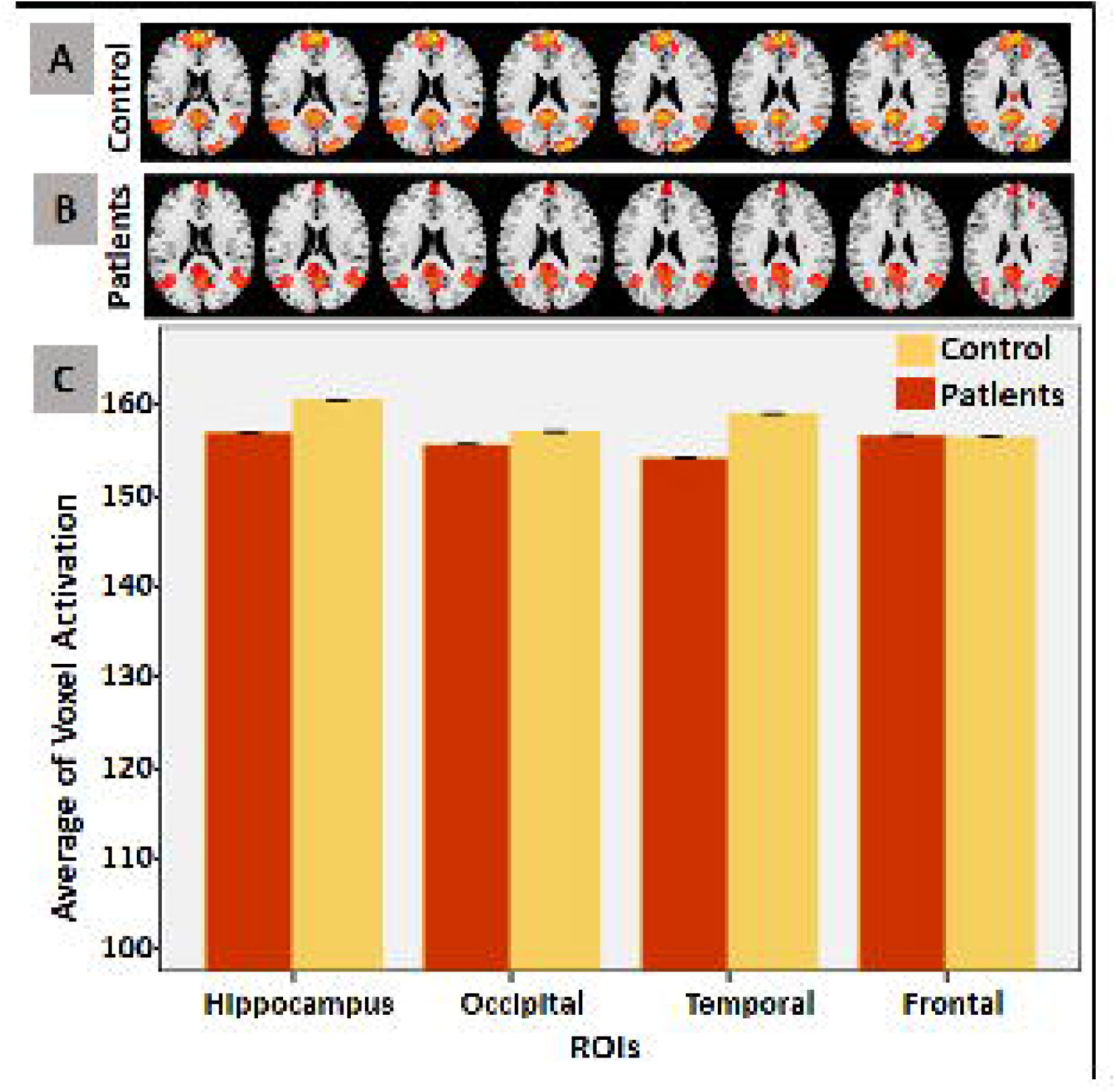
Brain Activation in Schizophrenia Patients in Comparison with Healthy Individuals. Axial slices of activations in HC, TL, OL, and FL of: **(A)** healthy individuals (control), and **(B)** schizophrenia patients, at resting state. **(C)** Bar chart of average activations in ROIs for control and patients.

### 3.2. Functional connectivity among the HC, OL, TL, and FL differs significantly in patients with SZ

The human brain is a complex neuronal network with variable levels of FC among different regions. FC gives rise to the physiological functions of different regions of the brain. Having observed significant variation in the activation of the four ROIs, we then evaluated the effect of this activation on the FC among these ROIs (28). To do this, we quantified the FC among the four ROIs by computing the Pearson correlations for the SZ and control groups (Figures 2A, B). For the controls, the highest positive correlation was observed between the FL and the TL (*r* = 0.85, *p* < 0.05) while the lowest positive correlation was between the FL and the OL (*r* = 0.63, *p* < 0.05) (Figure 2A). The smallest negative correlation in the controls was observed between the HC and the TL (*r* = −0.06), while the largest negative correlation was between the HC and the OL (*r* = −0.01). For the SZ groups, the highest positive correlation was observed between the FL and the TL (*r* = 0.79, *p* < 0.05), while the lowest positive correlation was between the FL and the OL (*r* = −0.26) (Figure 2B). The smallest negative correlation was between the HC and the TL (*r* = 0.17, p < 0.05), while the largest negative correlation was between the HC and the OL (*r* = 0.42, p < 0.05). These results indicate that the functional connectivity network is significantly altered in patients with SZ (Figures 2C, D).

**Figure 2.**
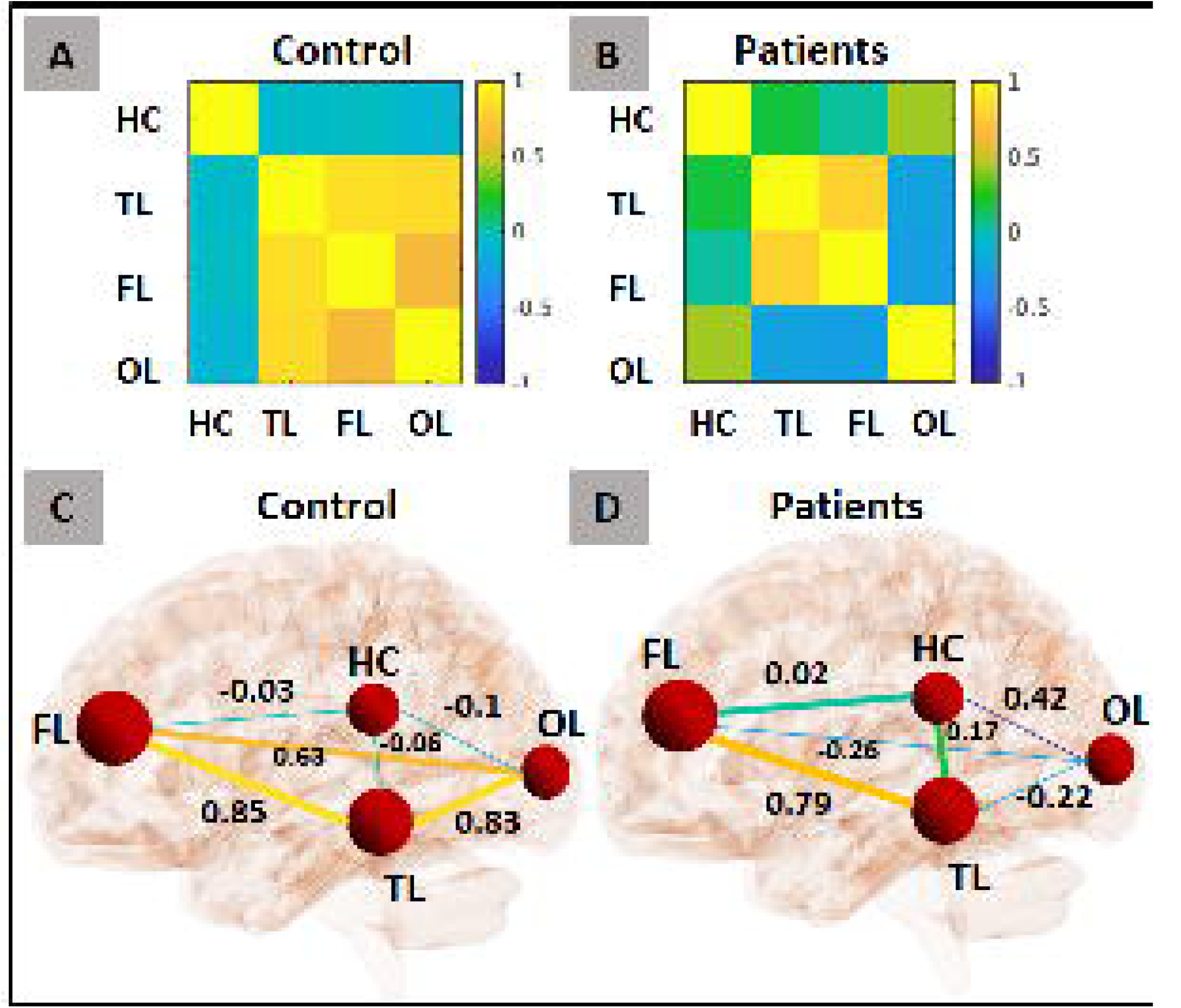
Correlation Between ROIs in Schizophrenia. Person correlation between ROIs in **(A)** Control, **(B)** Schizophrenia patients. Connectivity in ROIs in **(C)** Control, and **(D)** Schizophrenia patients, using BrainNet Viewer (52).

### 3.3 Patients with SZ exhibit aberrant structural connectivity in the ROIs

Next, to investigate the temporal variations in FC patterns (or *effective structural connectivity*) among the four ROIs, we employed the vector auto regression (VAR) model (51) at times *t*-2, *t*-1, and *t*. Between *t*-1 and *t*, the HC exhibited strong causal connectivity with the 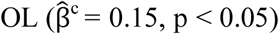 and with the 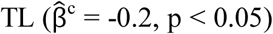 in healthy individuals. In patients with SZ, these connections of the HC with the OL and the TL were completely inhibited (Figure 3; Tables 1, 2). The TL also had strong causal connectivity with the 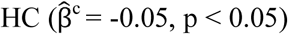 and the 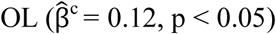 in the controls; these connections were also deregulated in the SZ group. The OL had causal connectivity with the 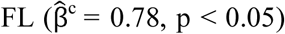 and vice versa 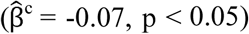, and the OL had causal connectivity with the 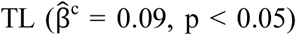 in the controls; these connections did not appear in patients with SZ. Furthermore, at lag times *t*-1 and *t*-2, the HC exhibited significantly diminished FC in patients with SZ. This differed from the controls most remarkably at lag *t*-2. Interestingly, at *t*-1 and *t*-2, the FL and OL showed enhanced anomalous connectivity in patients with SZ. The TL maintained its effective structural connectivity in both the controls and the patients with SZ at lag *t*-2.

**Table 1.** Estimated VAR Model for Control. **“***Mod (1-4) at lag1*” is the VAR model till lag 1, 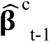 is the coefficient of variation for control case at lag 1, 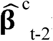 is coefficient of variation for control case at lag 2, *t* is the test statistic, S.E is the standard error of the model, *p* value is the probability of obtaining test results by chance.

| Mod<br>(1-4) at<br>lag1 | $\hat{\beta}_{t-1}^c$ | S.E | t | p | Mod<br>(1-4), at<br>lag 2 | $\hat{\beta}_{t-2}^c$ | S.E | t | p |
| --- | --- | --- | --- | --- | --- | --- | --- | --- | --- |
| HC(t) ← HC (t-1) | 0.51 | 0.05 | 10.15 | 0.000 | HC(t-2) | 0.17 | 0.05 | 3.4 | 0.001 |
| HC(t) ← OL (t-1) | 0.15 | 0.07 | 2.14 | 0.032 | OL(t-2) | -0.17 | 0.07 | -2.5 | 0.011 |
| HC(t) ← FL (t-1) | -0.00 | 0.09 | -0.08 | 0.937 | FL(t-2) | 0.03 | 0.09 | 0.3 | 0.714 |
| HC(t) ← TL (t-1) | -0.20 | 0.07 | -2.62 | 0.009 | TL(t-2) | 0.16 | 0.07 | 2.1 | 0.036 |
| OL(t) ← HC (t-1) | -0.05 | 0.03 | -1.35 | 0.178 | HC(t-2) | -0.02 | 0.03 | -0.7 | 0.478 |
| OL(t) ← OL (t-1) | 0.73 | 0.05 | 13.61 | 0.000 | OL(t-2) | 0.01 | 0.05 | 0.2 | 0.832 |
| OL(t) ← FL (t-1) | 0.22 | 0.07 | 3.16 | 0.002 | FL(t-2) | -0.11 | 0.07 | -1.6 | 0.098 |
| OL(t) ← TL (t-1) | 0.04 | 0.05 | 0.75 | 0.456 | TL(t-2) | 0.04 | 0.06 | 0.7 | 0.453 |
| FL(t) ← HC (t-1) | -0.01 | 0.03 | -0.60 | 0.547 | HC(t-2) | -0.00 | 0.03 | -0.2 | 0.777 |
| FL(t) ← OL(t-1) | -0.07 | 0.04 | -1.66 | 0.098 | OL(t-2) | 0.05 | 0.04 | 1.1 | 0.234 |
| FL(t) ← FL (t-1) | 0.78 | 0.05 | 14.15 | 0.000 | FL(t-2) | 0.03 | 0.05 | 0.6 | 0.495 |
| FL(t) ← TL (t-1) | 0.09 | 0.04 | 2.02 | 0.043 | TL(t-2) | 0.01 | 0.04 | 0.4 | 0.681 |
| TL(t) ← HC (t-1) | -0.05 | 0.03 | -1.66 | 0.097 | HC(t-2) | 0.01 | 0.03 | 0.4 | 0.644 |
| TL(t) ← OL (t-1) | 0.12 | 0.04 | 2.58 | 0.010 | OL(t-2) | -0.11 | 0.04 | -2.3 | 0.020 |
| TL(t) ← FL (t-1) | 0.07 | 0.06 | 1.15 | 0.251 | FL(t-2) | 0.03 | 0.06 | 0.6 | 0.541 |
| TL(t) ← TL (t-1) | 0.67 | 0.05 | 12.46 | 0.000 | TL(t-2) | 0.11 | 0.05 | 2.1 | 0.035 |

**Table 2.** Estimated VAR Model for Schizophrenia Patients. **“***Mod (1-4) at lag1*” is the VAR model till lag 1, 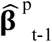 is the coefficient of variation for patients at lag 1, 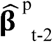 is coefficient of variation for patients at lag 2, *t* is the test statistic, S.E is the standard error of the model, *p* value is the probability of obtaining test results by chance.

| Mod<br>(1-4) at<br>lag1 | $\hat{\beta}_{t-1}^p$ | S.E | t | p | Mod<br>(1-4), at<br>lag 2 | $\hat{\beta}_{t-2}^p$ | S.E | t | p |
| --- | --- | --- | --- | --- | --- | --- | --- | --- | --- |
| HC(t) ← HC (t-1) | 0.36 | 0.05 | 6.34 | 0.000 | HC(t-2) | 0.29 | 0.05 | 5.0 | 0.000 |
| HC(t) ← OL (t-1) | 0.10 | 0.06 | 1.69 | 0.091 | OL(t-2) | -0.01 | 0.06 | -0.2 | 0.820 |
| HC(t) ← FL (t-1) | -0.00 | 0.07 | -0.10 | 0.917 | FL(t-2) | 0.03 | 0.07 | 0.4 | 0.638 |
| HC(t) ← TL (t-1) | 0.10 | 0.07 | 1.38 | 0.166 | TL(t-2) | -0.11 | 0.07 | -1.5 | 0.129 |
| OL(t) ← HC (t-1) | 0.09 | 0.04 | 1.96 | 0.050 | HC(t-2) | -0.10 | 0.04 | -2.04 | 0.042 |
| OL(t) ← OL (t-1) | 0.62 | 0.05 | 11.8 | 0.000 | OL(t-2) | 0.07 | 0.05 | 1.45 | 0.148 |
| OL(t) ← FL (t-1) | -0.01 | 0.06 | -0.22 | 0.826 | FL(t-2) | 0.03 | 0.06 | 0.58 | 0.562 |
| OL(t) ← TL (t-1) | 0.02 | 0.06 | 0.33 | 0.745 | TL(t-2) | 0.10 | 0.06 | 1.70 | 0.089 |
| FL(t) ← HC (t-1) | -0.03 | 0.04 | -0.65 | 0.513 | HC(t-2) | 0.05 | 0.04 | 1.14 | 0.256 |
| FL(t) ← OL (t-1) | 0.13 | 0.05 | 2.65 | 0.008 | OL(t-2) | -0.02 | 0.05 | -0.40 | 0.693 |
| FL(t) ← FL (t-1) | 0.31 | 0.06 | 5.10 | 0.000 | FL(t-2) | 0.25 | 0.06 | 4.21 | 0.000 |
| FL(t) ← TL (t-1) | 0.14 | 0.06 | 2.39 | 0.017 | TL(t-2) | -0.01 | 0.06 | -0.27 | 0.789 |
| TL(t) ← HC (t-1) | -0.06 | 0.04 | -1.47 | 0.142 | HC(t-2) | -0.01 | 0.04 | -0.30 | 0.768 |
| TL(t) ← OL (t-1) | 0.03 | 0.04 | 0.76 | 0.449 | OL(t-2) | 0.09 | 0.04 | 1.93 | 0.053 |
| TL(t) ← FL (t-1) | 0.02 | 0.05 | 0.46 | 0.073 | FL(t-2) | -0.06 | 0.05 | -1.13 | 0.259 |
| TL(t) ← TL (t-1) | 0.69 | 0.05 | 12.3 | 0.000 | TL(t-2) | 0.21 | 0.05 | 3.86 | 0.000 |

**Figure 3.**
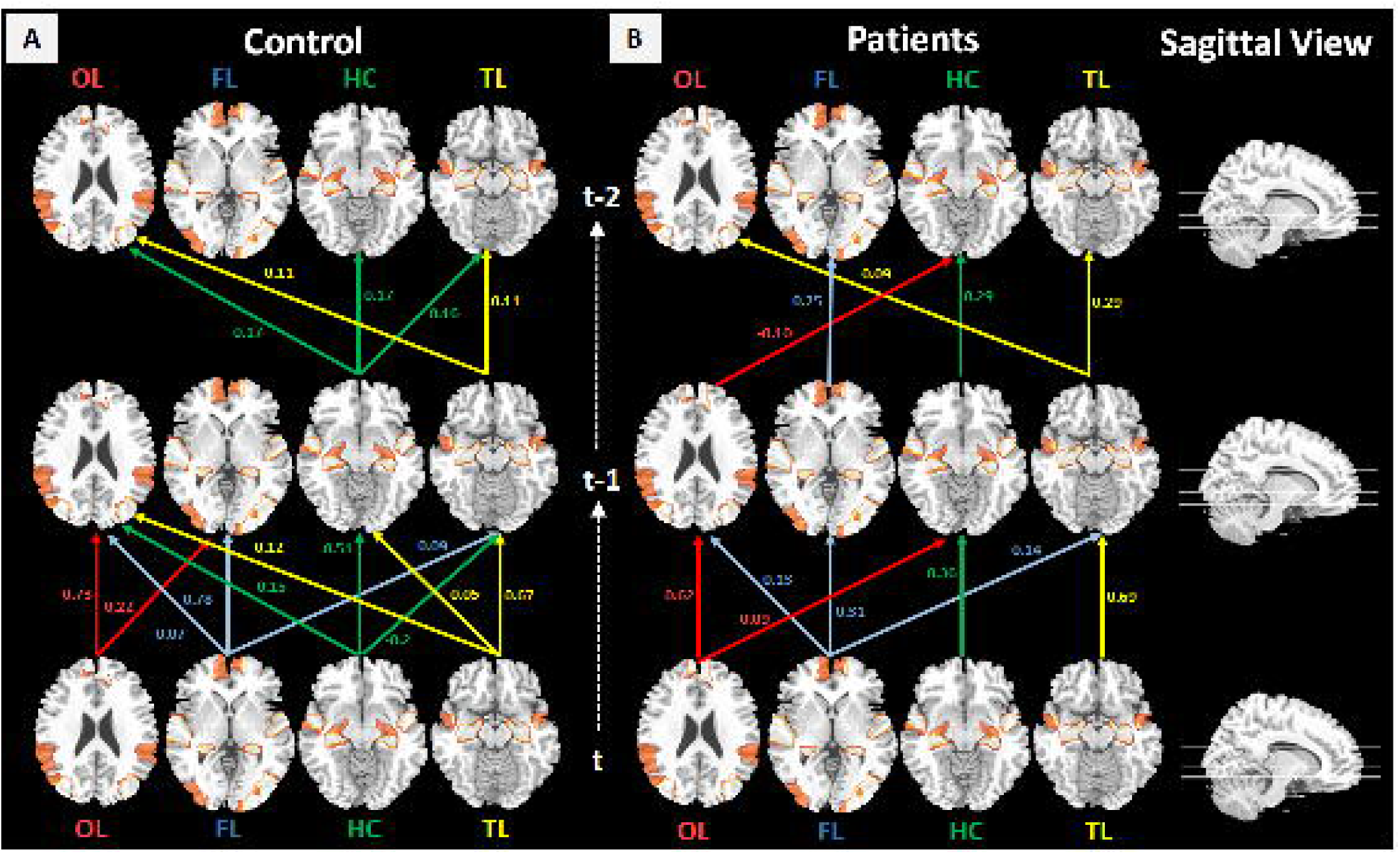
Structural Connectivity in ROIs. **(A)** Significant causality links in control between HC, OL, FL and TL, **(B)** Significant causality links in patients between HC, OL, FL and TL.

### 3.4 Validation of VAR models using Granger causality

To validate the structural model of the brain connectivity network obtained through VAR, we further employed a G-causal test (44) to examine the bidirectional causal effects of the ROIs on each other in patients with SZ and in the controls. In the controls (Table 3) the OL and TL regions had significant G-causality with the HC (Figure 4A). Moreover, the HC and the FL were G-causal for OL activation (Figure 4B), while the OL and the FL had G-causality with the TL (Figure 4C). In the controls, the TL also had G-causality with the FL (Figure 4D). However, we observed weaker causal relations among the four ROIs in patients with SZ (Table 4) than in the controls. Specifically, the HC was not G-causal for the other ROIs in patients with SZ (Figure 4A). However, the TL and the HC continued to have a significant G-causal relationship with the OL (Figure 4B). Moreover, the OL was G-causal for the TL (Figure 4C), while the OL and the TL were G-causal for the FL (Figure 4D). Taken together, these findings confirm our earlier hypotheses on aberrant patterns and reduced activation in the four ROIs in patients with SZ.

**Table 3:** Granger Causality Wald Test for Control. Equation is a dependent variable while excluded is representing independent variable,chi^2^ is a test of association, df stands for degree of freedom, Prob>chi^2^ is the probability value for drawing conclusion about null hypothesis.

| Equation | Excluded | Chi<sup>2</sup> | df | Prob>chi<sup>2</sup> |
| --- | --- | --- | --- | --- |
| HC | OL | 6.628 | 2 | 0.036 |
|  | FL | 0.223 | 2 | 0.894 |
|  | TL | 6.970 | 2 | 0.031 |
|  | ALL | 13.056 | 6 | 0.042 |
| OL | HC | 6.265 | 2 | 0.044 |
|  | FL | 11.206 | 2 | 0.004 |
|  | TL | 3.815 | 2 | 0.148 |
|  | ALL | 30.024 | 6 | 0.000 |
| FL | HC | 1.171 | 2 | 0.557 |
|  | OL | 2.751 | 2 | 0.253 |
|  | TL | 10.839 | 2 | 0.004 |
|  | ALL | 15.068 | 6 | 0.020 |
| TL | HC | 3.450 | 2 | 0.178 |
|  | OL | 7.001 | 2 | 0.030 |
|  | FL | 6.353 | 2 | 0.042 |
|  | ALL | 19.55 | 6 | 0.003 |

**Table 4.**
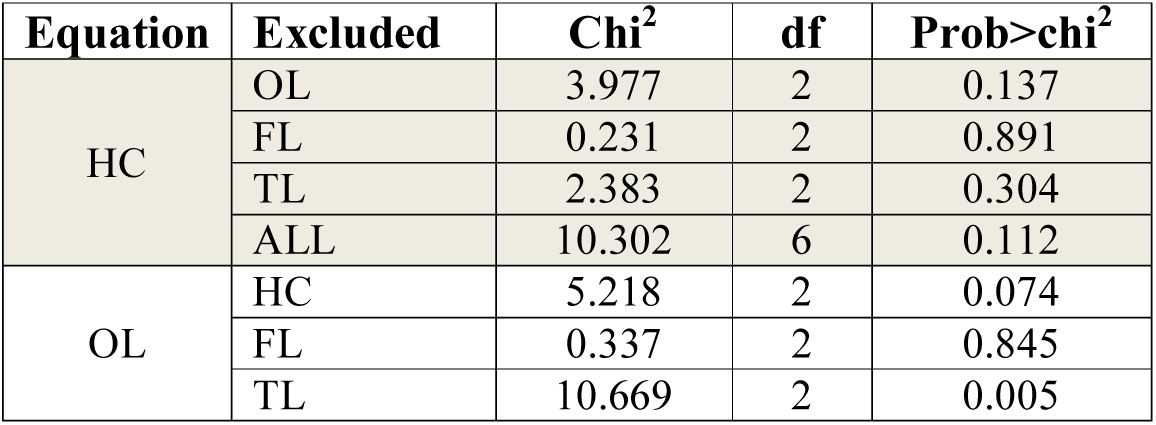
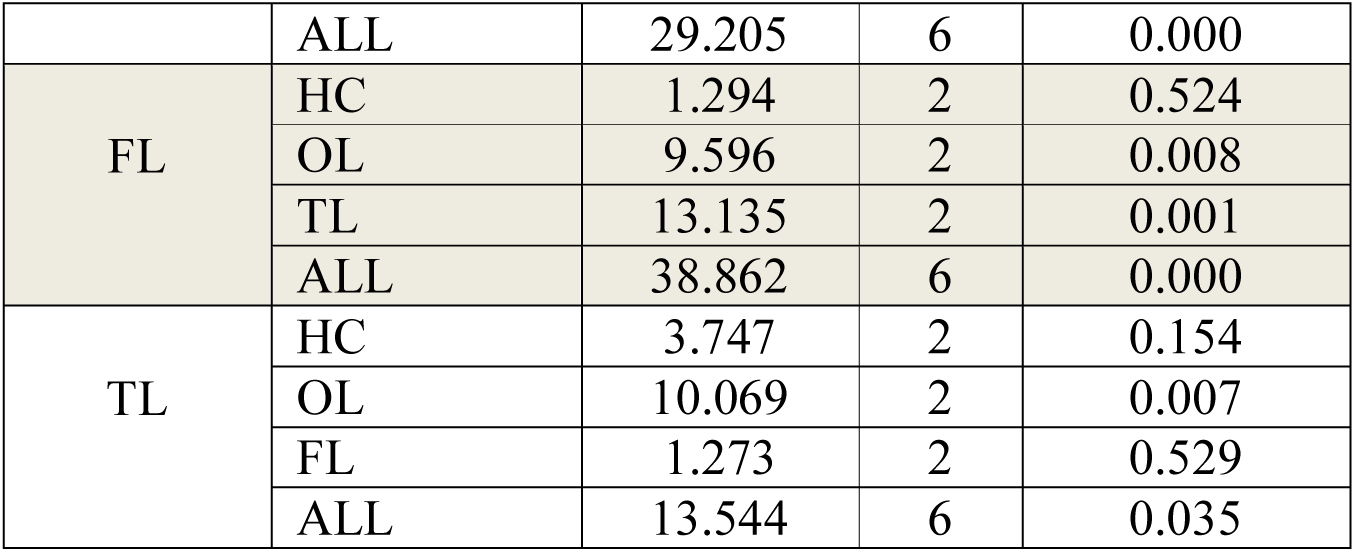
Granger Causality Wald Test for Patients. Equation is a dependent variable while excluded is representing independent variable chi^2^ is a test of association, df stands for degree of freedom, Prob>chi^2^ is the probability value for drawing conclusion about null hypothesis.

**Figure 4.**
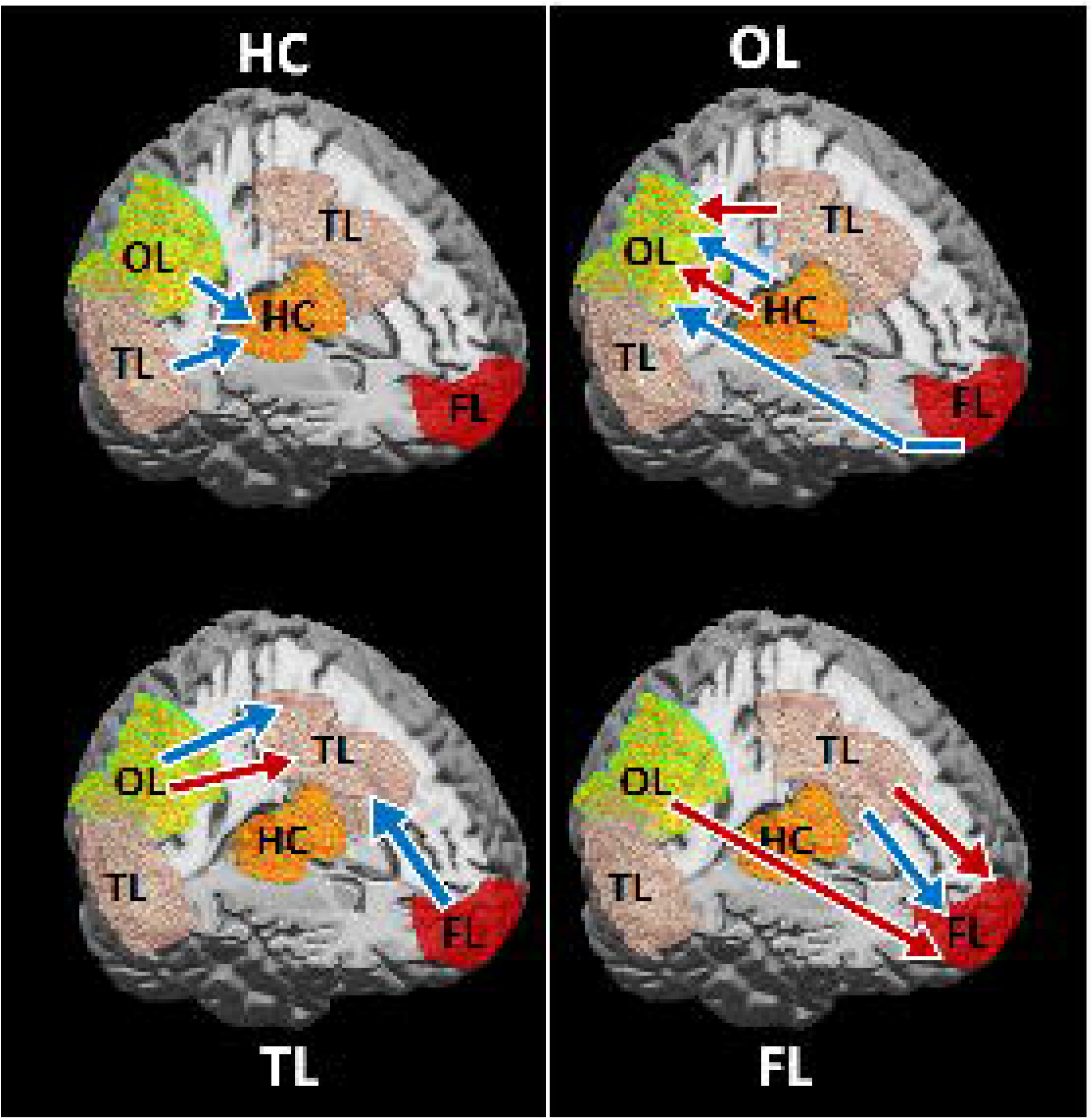
Structural Connectivity in ROIs obtained through Granger Causality for HC, OL, TL, and FL. Blue and Red arrows show significant granger causality between ROIs for the control case and SZ patients, respectively.

### 3.5 Causal analysis through mediation reveals differences in decision making in patients with SZ

To investigate differences in the decision-making processes of patients with SZ, we performed a causal analysis of HC mediation. To do this, we analyzed the role of the HC as a mediator between the FL and OL in the controls and in patients with SZ (Figure 5). Our results show that the HC played a significant role in the patients with SZ (Figure 5B) but was non-significant in the control group (Figure 5A). We further observed that the FL disturbed the activity of the OL and the HC in patients with SZ, while in the controls this effect was non-significant (Table 5).

**Table 5.** Mediation Analysis for Regions of Interest. S.E. stands for standard error, C.R. is the confidence ratio and P is the probability value.

|  |  |  |  |  |  |  |  |
| --- | --- | --- | --- | --- | --- | --- | --- |
| Control | Dependent | Effect | Independent | Estimate | S.E. | C.R. | P |
|  | HC | <--- | OL | -0.048 | 0.045 | -1.062 | 0.288 |
|  | FL | <--- | OL | 0.476 | 0.035 | 13.791 | *** |
|  | FL | <--- | HC | 0.225 | 0.038 | 5.904 | *** |
| Patients | Dependent | Effect | Independent | Estimate | S.E. | C.R. | P |
|  | HC | <--- | OL | 0.39 | 0.038 | 10.333 | *** |
|  | FL | <--- | OL | 0.464 | 0.035 | 13.098 | *** |
|  | FL | <--- | HC | 0.361 | 0.042 | 8.676 | *** |

**Figure 5.**
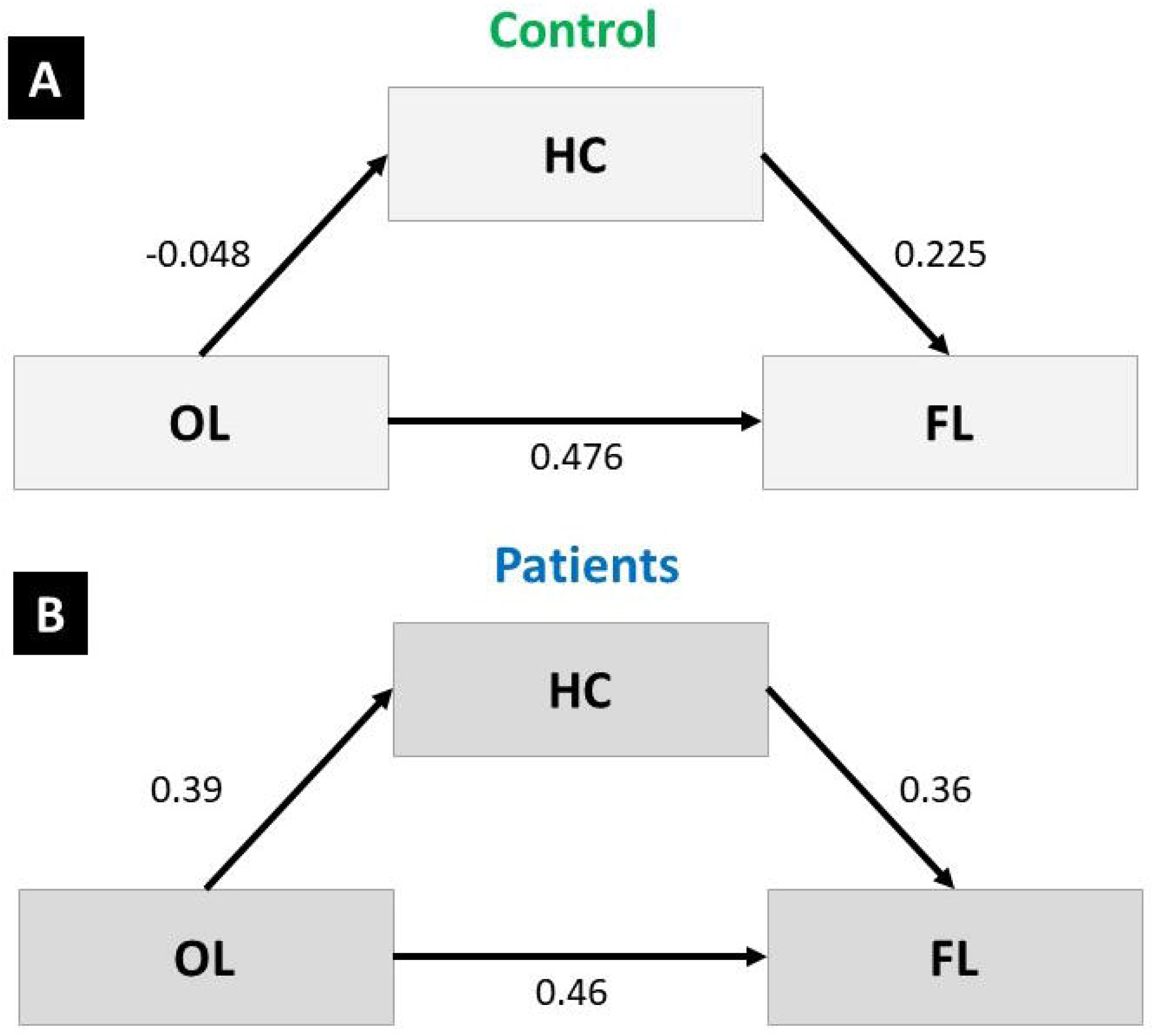
Mediation Analysis in ROIs. **(A)** Significant mediation in control between HC, OL, and FL, **(B)** Significant mediation in patients between HC, OL, and FL.

## 4. DISCUSSION

This study examined rsFC in patients with SZ on a whole-brain scale. The results show that the parietal region has less connectivity to bilateral DLPFC, while the parietal and frontal regions have strong connectivity in patients with SZ. Our results also showed that patients with SZ exhibited significantly less activation in the HC, the OL, and the TL than healthy controls (61), while the FL showed significantly more activation in patients with SZ. Abnormal functional connectivity in the prefrontal cortex may reflect psychopathologies such as an inability to allocate internal or external attentional resources, a crucial skill for goal-oriented behaviors (61, 62). Working memory and decision-making deficits have also been repeatedly reported in patients with SZ; these symptoms are often linked to abnormal functioning of the prefrontal cortex (63, 64). Temporal hallucinations and delusions are the main characteristics of SZ; these are mainly attributed to aberrant FC in the temporal cortex (65, 66). There is also a well-established link between SZ and the hippocampus, a complex region of the brain that plays a critical role in multiple cognitive domains, including memory, imagination, and emotions (67, 68), that are known to be impaired in patients with SZ (69).

Approaches such as spatial independent component analysis (ICA), seed-based temporal correlation analysis (SCA) and Granger causality analysis (GCA) can be used to explore the activity and functions of the intrinsic neuronal network using rsfMRI data (70–73). Our findings are consistent with those of other studies. For example, one study by (74) observed reduced degree centrality (DC) of the bilateral putamen nuclei in patients with SZ compared to controls. DC is an index used to identify the regions of the brain (at the whole-brain level) that display functional deficits in patients with SZ. That study also observed a lack of causal connectivity between the putamen and multiple regions of the default mode network (DMN), the orbital area of the inferior frontal cortex, and the right fusiform in patients with SZ. In addition, a previous study found abnormal rsFC in the amygdala subregional-sensorimotor regions of the brain in patients with SZ; this abnormality was also associated with positive symptoms in patients with SZ (75). A similar pattern of altered rsFC has been observed throughout the entire brain in patients with SZ. Disrupted pathways from the limbic areas to the thalamus have also been demonstrated using resting-state effective connectivity (rsEC) analysis (76).

Furthermore, a wide range of alterations to thalamic nuclei functional connectivity have been observed in the cortico-cerebellar-thalamo-cortical circuit pathways of patients with SZ (77). A systematic review of task and rsfMRI studies demonstrated the convergence of brain neural dysfunction between tasks and rsfMRI abnormalities in the prefrontal regions, including the dorsal lateral prefrontal cortex, the orbital frontal cortex, and the TL, particularly the superior temporal gyrus (78). Together, these previous findings demonstrate that patients with SZ exhibit altered FC and effective connectivity (EC) among large regions of the brain. Moreover, brain rsFC and rsEC can be used as diagnostic markers for SZ and might be implicated in therapeutic interventions as well.

A major strength of this study is its use of a combination of three different FC analysis methods to investigate functional disconnections between various regions of the brain. However, the study does have some limitations, which should be considered when interpreting the results. Most significantly, the sample size in the present study is small. Additional longitudinal follow-up studies with larger samples are needed to elucidate the alterations in FC between regions of the brain in patients with SZ.

## Conclusion

GCA is a useful tool for characterizing the functional direction of time-series data. GCA has broad implications in the neurosciences and neuroimaging because of the importance of the FC of different regions of the brain during tasks. GCA can be used to characterize the significant functional directions of different regions of the brain. In the above diagram, the functional direction of different regions of the brain are indicated by arrows. These arrows illustrate the significant Granger causes in the patients with SZ (red) and the control subjects (black) in our study. We can conclude that, in patients with SZ, some regions of the brain are less active than those of healthy subjects during task performance.

## Ethical publication statement

We confirm that we have read the Journal’s position on issues involved in ethical publication and affirm that this report is consistent with those guidelines.

## Acknowledgements

The authors would like to thank participants and their family members.

